# *Plasmodium falciparum* CRK4 links early mitotic events to the onset of S-phase during schizogony

**DOI:** 10.1101/2022.08.31.505163

**Authors:** Marta Machado, Severina Klaus, Darius Klaschka, Julien Guizetti, Markus Ganter

**Author notes:** **Correspondence:** Markus Ganter.

## Abstract

*Plasmodium falciparum* proliferates through schizogony in the clinically relevant blood stage of infection. During schizogony, consecutive rounds of DNA replication and nuclear division give rise to multinucleated stages before cellularization occurs. Although these nuclei reside in a shared cytoplasm, DNA replication and nuclear division occur asynchronously. Here, by mapping the proteomic context of the S-phase-promoting kinase *Pf*CRK4, we show that it has a dual role for nuclear-cycle progression: *Pf*CRK4 orchestrates not only DNA replication, but also the rearrangement of intranuclear microtubules. Live-cell imaging of a reporter parasite showed that these microtubule rearrangements coincide with the onset of DNA replication. Together, our data render *Pf*CRK4 the key factor for nuclear-cycle progression, linking entry into S-phase with the initiation of mitotic events. In part, such links may compensate for the absence of canonical cell cycle checkpoints in *P. falciparum*.

## Introduction

*P. falciparum* is the etiologic agent of the most severe form of human malaria, which remains a major cause of global morbidity and mortality (WHO, 2021). In the clinically relevant blood stage of infection, this unicellular eukaryotic parasite proliferates through a diverged cell cycle mode called schizogony, where alternating rounds of S-phase and nuclear division form multinucleated cells before cellularization occurs (Klaus et al., 2022; Rudlaff et al., 2020). In contrast to other multinucleated cells, e.g., the early *Drosophila* embryo (Farrell and O’Farrell, 2014), *P. falciparum* nuclei undergo DNA replication and nuclear divisions asynchronously despite sharing a common cytoplasm (Klaus et al., 2022; McDonald and Merrick, 2022; Read et al., 1993; Simon et al., 2021). This suggests a divergent and locally restricted regulation of nuclear cycle progression (Francia and Striepen, 2014; Gubbels et al., 2021; Matthews et al., 2018).

In other eukaryotes, cyclins and cyclin-dependent kinases (CDKs) drive the timely progression through the cell cycle (Basu et al., 2022; Malumbres, 2014). While no canonical G1, S- and M-phase cyclins could be identified (Robbins et al., 2017; Roques et al., 2015), the *P. falciparum* genome harbors CDK-like kinases and several related kinases, such as the cdc2-related kinase 4 (*Pf*CRK4) (Doerig et al., 2002). Conditional depletion of *Pf*CRK4 suppressed DNA replication in the blood stage of infection and this kinase is essential for the initial and subsequent rounds of nuclear multiplication (Ganter et al., 2017). Phosphoproteomic profiling of *Pf*CRK4 identified a set of potential effector proteins, which are likely involved in origin of replication firing (Ganter et al., 2017). This suggested that *Pf*CRK4 is a major S-phase promoting factor in *P. falciparum*. However, a detailed understanding of the regulatory circuits that drive asynchronous nuclear cycles during blood-stage schizogony is still missing.

To gain a better understanding of *Pf*CRK4’s role for asynchronous nuclear cycles, we profiled the proteomic context of this kinase and found several microtubule-associated proteins in the vicinity of *Pf*CRK4, suggesting an additional molecular function besides regulating DNA replication. Further supporting this notion, we detected *Pf*CRK4 predominantly at intranuclear microtubule foci. Super-resolution live-cell imaging showed that major microtubule rearrangements depend on *Pf*CRK4 activity and coincide with the onset of DNA replication. Our data integrates the dynamics of microtubule structures and DNA replication, which have been independently used to assess nuclear cycle progression in *P. falciparum*. These data also expand the repertoire of molecular functions assigned to *Pf*CRK4, rendering it a key regulator of nuclear-cycle progression during asynchronous nuclear cycles.

## Results

To analyze the role of the S-phase promoting kinase *Pf*CRK4 for nuclear-cycle progression, we mapped its proteomic context. For this, we endogenously fused *Pf*CRK4 with the promiscuous biotin ligase BirA*, which allows the biotinylation of proteins within a radius of ~20 nm (Roux et al., 2012) (Fig. 1A; S1A, B). To functionally characterize the *Pf*CRK4::BirA* expressing line, we stained parasites with fluorescent streptavidin to detect biotinylated proteins (Fig. 1B). Wild type (WT) parasites cultures supplemented with 50 μM biotin showed no detectable signal. In *Pf*CRK4::BirA* parasites, the approximately 0.8 μM biotin of the standard culture medium was sufficient for a detectable biotin labeling, as previously reported (Boucher et al., 2018; Khosh-Naucke et al., 2018). The signal strongly increased, when we supplemented the culture media with 50 μM biotin and we found biotinylated proteins predominantly localized to nucleus of *Pf*CRK4::BirA* parasites (Fig. 1B, S1C). This is in agreement with the spatiotemporal expression of *Pf*CRK4 fused to GFP (Fig. S1A, B, D) and the previously reported localization of *Pf*CRK4 (Ganter et al., 2017). Western blot analysis further supported these findings (Fig. 1C), indicating that the *Pf*CRK4::BirA* fusion protein is active.

**Fig. 1.**
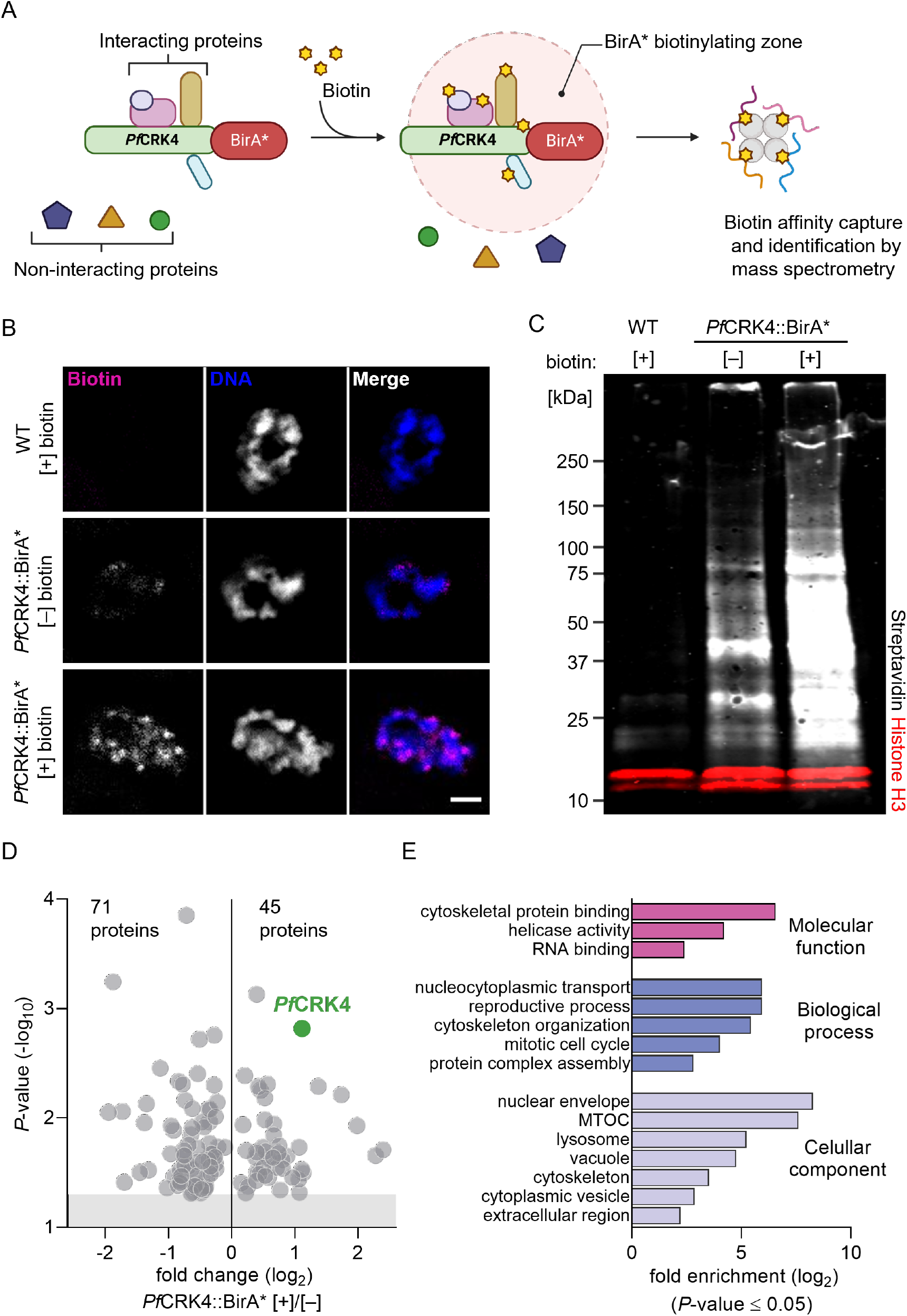
Profiling the proximity proteome of *Pf*CRK4 identified a potential link to microtubules. **A** Scheme illustrating the principal steps of proximity-based labeling via BioID, created with BioRender.com. **B** Biotinylated proteins predominantly localize to a small region of *P. falciparum* nuclei; scale bar, 2 μm. **C** Western blot analysis showed specific biotinylation in *Pf*CRK4::BirA* parasites. IRDye 800CW streptavidin was used to detect biotinylated proteins (grey), a-histone H3 was used as a loading control and visualized by IRDye 680CW Goat anti-rabbit IgG (red). **D** Under- and overrepresented proteins in the vicinity of *Pf*CRK4::BirA* parasites [+] 50 μM biotin relative to [−] biotin. Shown are mean values of triplicates; grey shaded region indicates a *P*-value ≥ 0.05 (Student’s *t*-test, two-tailed, equal variance). **E** Gene ontology term enrichment analysis of 44 proteins (excluding *Pf*CRK4) that are overrepresented in the vicinity of *Pf*CRK4, calculated using the built-in tool at PlasmoDB.org.

To profile the proteomic context of *Pf*CRK4, we enriched for biotinylated proteins from *Pf*CRK4::BirA* [+] and [−] 50 μM biotin as well as WT parasites [+] 50 μM biotin and analyzed triplicate samples by quantitative label-free mass spectrometry. We refrained from a quantitative analysis of WT [+] 50 μM biotin samples, which were intended to serve as negative control, due to low reproducibility between triplicates (Fig. S2). Next, we asked which proteins were enriched in the proximity of *Pf*CRK4::BirA* by comparing the abundance of proteins in [+] over [−] biotin samples (Fig. 1D, Tab. S1). We found 44 proteins to be significantly overrepresented in the proximity of *Pf*CRK4, including the bait protein, and 71 proteins significantly underrepresented. The set of overrepresented proteins included four proteins that were also less phosphorylated in the absence of *Pf*CRK4 (Fig. S3, Tab. S1) (Ganter et al., 2017). To further validate our proximity proteome data, we assessed the localization of one of these candidates, the putative spindle pole body protein (*Pf*SPB, PF3D7_0303500) (Fig. S4). We detected *Pf*SPB at the parasite’s centrosome, also called centriolar plaque, which organizes intranuclear microtubules (Li et al., 2022; Rashpa and Brochet, 2022; Simon et al., 2021). At the centriolar plaque, *Pf*SPB localized between the extranuclear centrin and the intranuclear tubulin (Fig. S4C). This localization is consistent with the bulk labelling of biotinylated proteins and the localization of *Pf*CRK4::GFP (Fig. 1B; S1C, D).

Next, we used Gene Ontology (GO) term enrichment analysis to examine the functional characteristics of the under- and overrepresented sets of proteins in the proximity of *Pf*CRK4. While underrepresented proteins appear mainly involved in translation (Fig. S5A), the enriched GO terms of overrepresented proteins suggested an association of *Pf*CRK4 with the cytoskeleton (Fig. 1E). STRING analysis of the overrepresented proteins clustered *Pf*CRK4 with β-tubulin, suggesting a link between *Pf*CRK4 and microtubule structures of the cytoskeleton (Fig. S5B) (Szklarczyk et al., 2020). Together with the previous observation that *Pf*CRK4-depleted parasites show prominent intranuclear microtubule structures (Ganter et al., 2017), these data suggested that, besides directing DNA replication, *Pf*CRK4 possesses an additional role regulating microtubule dynamics during nuclear cycle progression.

Microtubules display a dynamic rearrangement during the nuclear cycle (Liffner and Absalon, 2021; Simon et al., 2021). Initially, microtubules form a so-called hemispindle, consisting of relatively long microtubules that radiate from the centriolar plaque into the nucleoplasm. Later they rearrange into short microtubules, which build the early mitotic spindle. The mature mitotic spindle eventually extends to partition the duplicated genome. After nuclear division, hemispindles re-appear in sister nuclei as the remnants of the divided anaphase spindle (Liffner and Absalon, 2021; Simon et al., 2021). To investigate the hypothesis that *Pf*CRK4 contributes to microtubule dynamics, we quantified its subcellular localization (Fig. 2; S6, S7). We detected *Pf*CRK4 predominantly in the nucleus and within the nuclei, the highest *Pf*CRK4 signal per area was observed at microtubule structures (Fig. 2A, B; S6). Comparing elongated with focal nuclear microtubule structures, we found *Pf*CRK4 predominantly associated with microtubule foci (Fig. 2A, C; S6). Together, these data further indicated a link between *Pf*CRK4 and intranuclear microtubule structures.

**Fig. 2.**
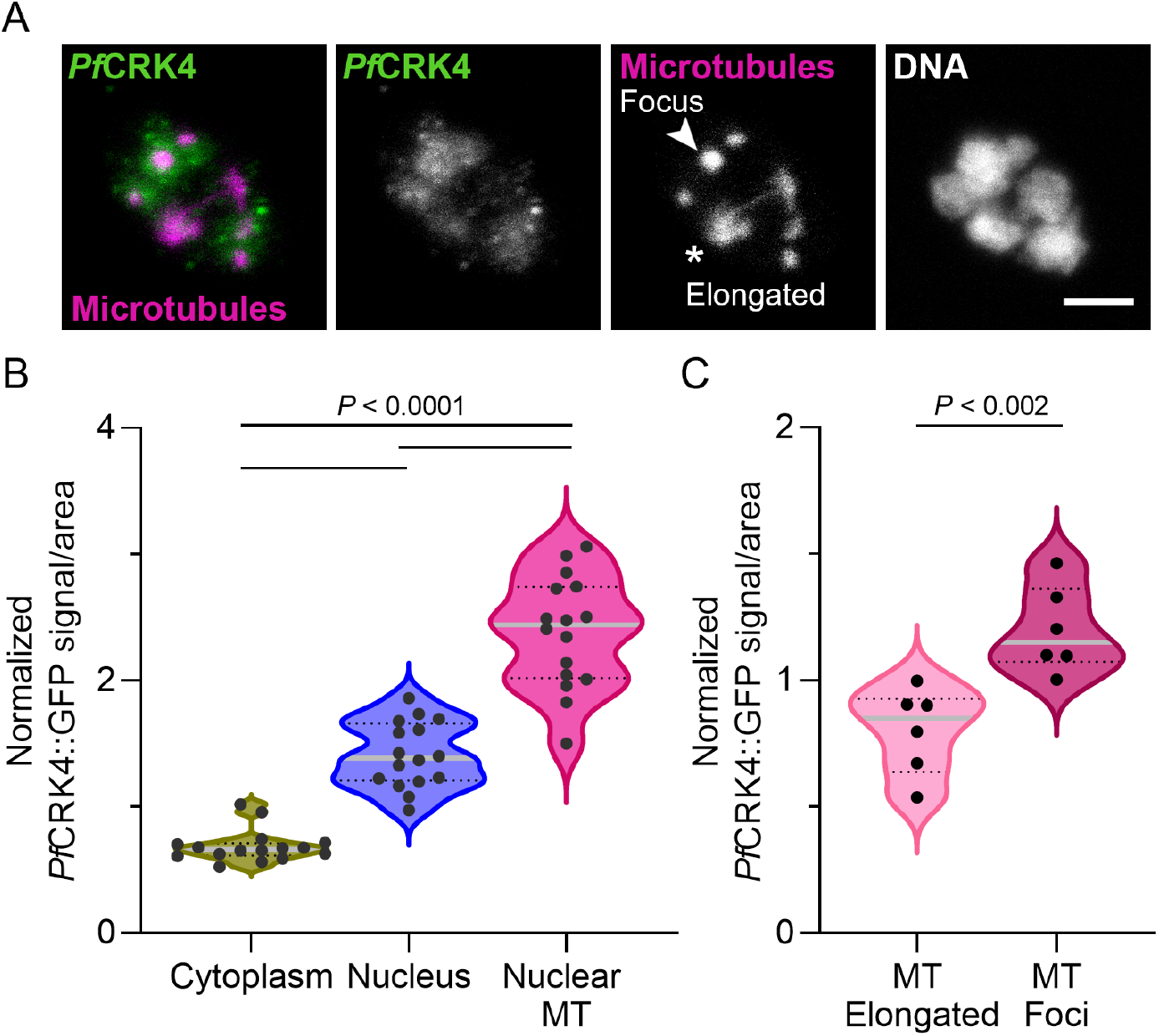
*Pf*CRK4 localized predominantly to focal microtubule structures in the nucleus. **A** Immunofluorescence showing uneven distribution of *Pf*CRK4 relative to nuclear microtubule structures in a multinucleated parasite; scale bar, 2 μm. **B** *Pf*CRK4 predominantly localizes to nuclear microtubule (MT) structures; *P*-value, one-way ANOVA with Tukey’s multiple comparison posttest. Signals were normalized to the total fluorescence of the whole cell. **C** *Pf*CRK4 associates more with focal than with elongated microtubule structures; *P*-value, unpaired Student’s *t*-test with Welch’s correction. Signals were normalized to the total fluorescence of nuclear microtubules.

To further explore this potential link, we employed the previously established *Pf*CRK4-HA-DD parasite, which permits the inducible depletion of *Pf*CRK4 by the removal of the small molecule Shield-1 (Ganter et al., 2017). First, we confirmed previous data that *Pf*CRK4-depleted parasites display prominent intranuclear microtubule structures that resemble hemispindles (Fig. 3A) (Ganter et al., 2017). Next, we analyzed the effect of restored *Pf*CRK4 activity on these microtubule structures. For this, we depleted *Pf*CRK4 until approximately 36 hours post invasion, when the *Pf*CRK4 phenotype is well-established (Ganter et al., 2017). Then, we either kept the kinase depleted or we rescued *Pf*CRK4 expression and, hence, kinase activity by addition of Shield-1 (Fig. 3B). Immediately after, we simultaneously analyzed the dynamics of intranuclear microtubules and the nuclear DNA content. For this, we stained the parasites with the live-cell compatible dyes SPY555-tubulin and 5’-SiR-Hoechst and imaged single cells at super-resolution for 4 hours.

**Fig. 3.**
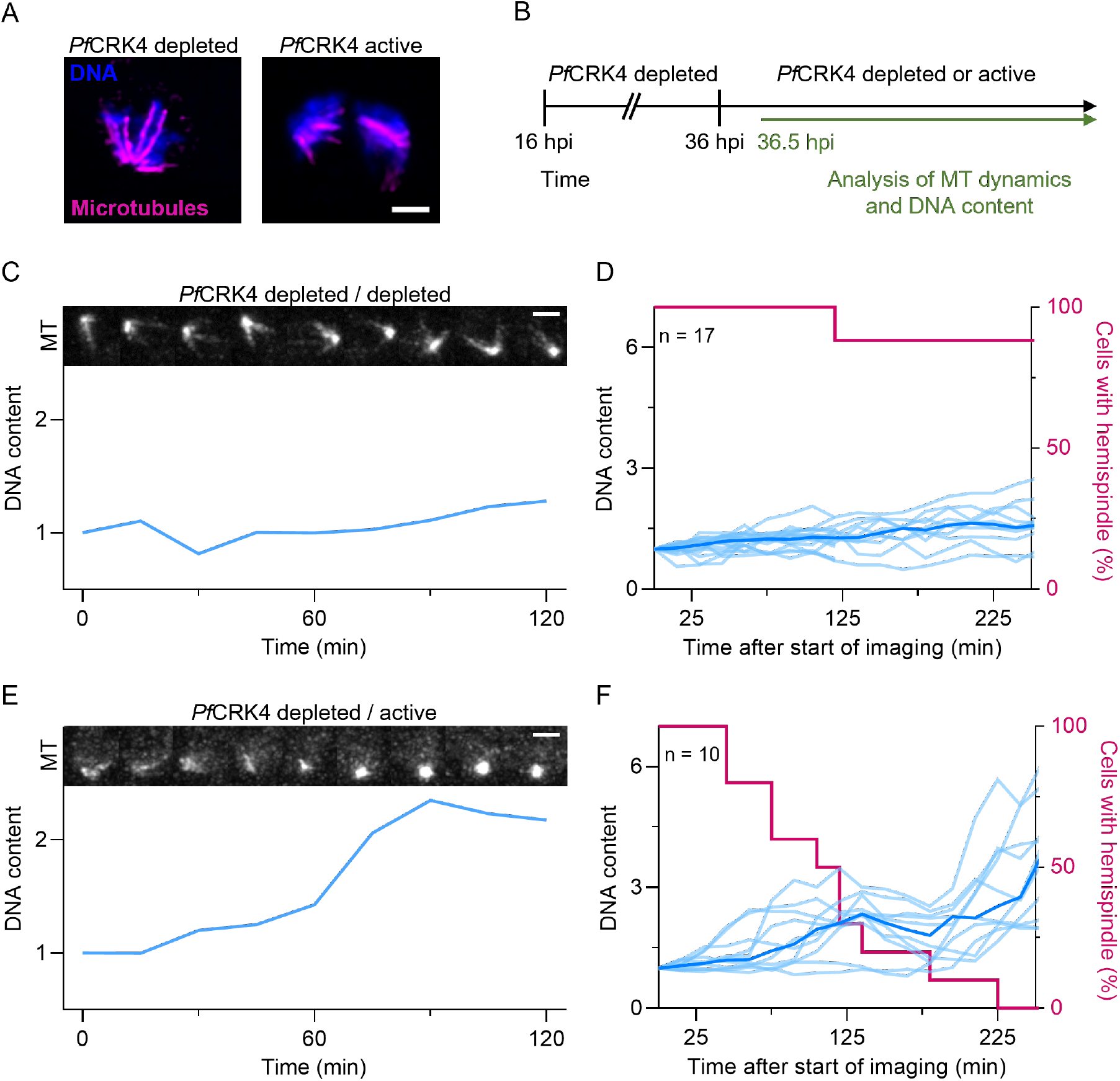
Rearrangement of nuclear microtubules depends on *Pf*CRK4 activity. **A** Immunofluorescence of nuclear microtubules in *Pf*CRK4-depleted and *Pf*CRK4-active parasites; scale bar, 1 μm. **B** Scheme of the timing of restored *Pf*CRK4 activity and subsequent super-resolution live-cell imaging; hpi, hours post invasion. **C** Representative images of microtubule (MT) structures (top) and normalized DNA content (bottom) over time in a *Pf*CRK4-depleted parasite, single z-slice, scale bar, 2 μm. **D** Quantification of hemispindle structures and the DNA content of *Pf*CRK4-depleted parasites; light blue, single cell signals; dark blue, mean signal. **E** Representative images of microtubule (MT) structures (top) and normalized DNA content (bottom) over time in a parasite with restored *Pf*CRK4 activity, single z-slice, scale bar, 2 μm. **F** Quantification of hemispindle structures and the DNA content of parasites with restored *Pf*CRK4 activity; light blue, single cell signals; dark blue, mean signal. C–F, DNA content was normalized to the total fluorescence of the first time point (36.5 hpi).

In *Pf*CRK4-depleted cells, we found prominent intranuclear microtubule structures, which consisted of constantly polymerizing and depolymerizing microtubules and a profoundly suppressed DNA replication (Fig. 3C, D; movies S1, S2). In contrast, when *Pf*CRK4 activity was rescued, the prominent hemispindle-like structures rearranged into focal microtubule structures and the DNA content of these nuclei increased (Fig. 3E, F; movies S3, S4), further coupling intranuclear microtubule dynamics to *Pf*CRK4 activity. To test whether microtubule rearrangements are a prerequisite for DNA replication, we treated the cells with the microtubule-stabilizing compound paclitaxel (Kumar, 1981) and then rescued *Pf*CRK4 activity. Next, we analyzed the microtubule structures by immunofluorescence and quantified the DNA content by flow cytometry (Ganter et al., 2017; Russo et al., 2009; Theron et al., 2010). Although we readily detected large aberrant microtubule structures in these cells, DNA replication commenced when *Pf*CRK4 was active (Fig. S8), suggesting that accurate microtubule rearrangement is not necessary for DNA replication to commence. Together, these data indicate that both rearrangement of intranuclear microtubules and DNA replication depend on *Pf*CRK4 activity.

As depletion of *Pf*CRK4 arrests normal nuclear cycle progression, we next investigated the temporal coordination of intranuclear microtubule rearrangement and DNA replication without perturbation. We stained microtubules with SPY555-tubulin in a nuclear cycle sensor line (Klaus et al., 2022). This *P. falciparum* line expresses the red-fluorescent protein mCherry in all nuclei and the proliferating cell nuclear antigen (PCNA) 1 fused to GFP. PCNA1::GFP transiently accumulates only in those nuclei that replicate their DNA, and hence, DNA replication and nuclear division events can be tracked. Using super-resolution live-cell microscopy, we imaged cells with a single nucleus and found that hemispindles with dynamic microtubules formed well before PCNA1::GFP accumulated in the nuclei, which marks the onset of DNA replication. Thus, the presence of a hemispindle is currently the first visible cue of an imminent S-phase. Coinciding with the start of DNA replication, hemispindles rearranged into short microtubule structures that likely localized to the centriolar plaque (Fig. 4A–C, movies S5, 6) (Simon et al., 2021). Once the subsequent nuclear division concluded, hemispindle structures re-appeared in sister nuclei. Again, the rearrangement of these second-generation hemispindles coincided with the onset of DNA replication in those nuclei (Fig. 4A–C; movies S5, S6), suggesting a tight temporal coordination of these two events throughout schizogony. We also found that PCNA1::GFP initially accumulated adjacent to the focal microtubule signal at the centriolar plaque and over time translocated to distal regions of the nucleus (Fig. 4A; movies S5, 6), suggesting that DNA close to the early mitotic spindle is replicated before DNA in the nuclear periphery, such as *var* genes (Lopez-Rubio et al., 2009). Throughout S-phase and after it concluded, the focal microtubule signals appeared to increase (Fig. 4A inset; movies S5, S6), potentially reflecting the maturation of the mitotic spindle.

**Fig. 4.**
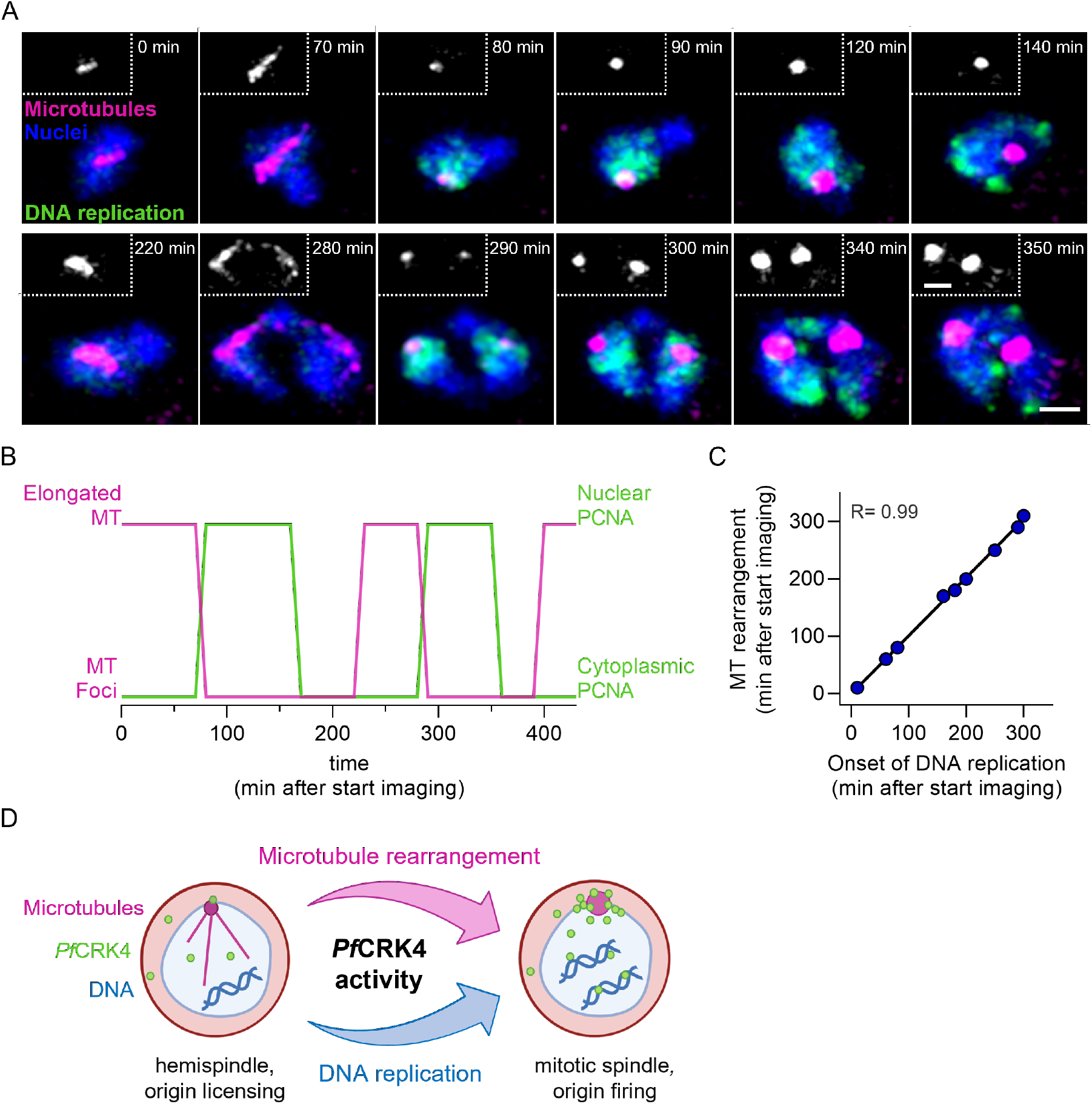
Rearrangement of nuclear microtubules coincides with the onset of S-phase. **A** Super-resolution live-cell imaging of microtubule structures (magenta) in a nuclear cycle sensor line, where nuclei are marked with mCherry (blue) and nuclear accumulation of PCNA1::GFP (green) marks DNA replication; inserts, microtubule signal; scale bars, 1 μm. **B** Timing of elongated and focal nuclear microtubule structures (MT) relative to the localization of PCNA1::GFP, with nuclear PCNA1::GFP marking DNA replication of the parasite shown in A. **C** Correlation of the timing of microtubule rearrangement and the onset of DNA replication. **D** Model of the localization and the key molecular functions of *Pf*CRK4, which drives microtubule rearrangements from hemispindle to early mitotic spindle and the onset of DNA replication.

Together, our data determined the temporal coordination of nuclear microtubule dynamics and DNA replication during nuclear cycle progression. We also found that both events depend on the activity of *Pf*CRK4, rendering this kinase a master regulator of nuclear cycle progression (Fig. 4D).

## Discussion

In the clinically relevant blood stage of infection, *P. falciparum* proliferates through schizogony, during which asynchronous nuclear cycles occur in a multinucleated cell. This mode of replication is *Plasmodium-specific* and, hence, a good target for therapeutic interventions. Yet, to exploit this potential, insights into the underlying molecular mechanisms are needed and we, therefore, further analyzed the role of the S-phase promoting kinase *Pf*CRK4 (Ganter et al., 2017). Combining proteomic and advanced imaging approaches, we provide evidence that intranuclear microtubule rearrange into early mitotic spindles at the onset of DNA replication, and that both processes are linked via *Pf*CRK4.

During normal nuclear cycle progression, hemispindles appear well before S-phase commences (Fig. 4A, B; movies S5, S6). Thus, in contrast to what has been proposed (Hawkins et al., 2022), the large hemispindle-like structures seen in *Pf*CRK4-depleted parasites are consistent with a developmental arrest just prior to S-phase, rather than representing a *Pf*CRK4-depletion phenotype during nuclear division (Fig. 3, 4). Whether the hemispindle has a biological function or whether it is a result of local high concentrations of tubulin that self-assemble into microtubules remains unclear (Simon et al., 2021). Similarly, the events that occur after the microtubule rearrange and that eventually lead to the attachment of the duplicated chromosomes to the mature mitotic spindle are not understood.

The molecular details of how *Pf*CRK4 regulates the rearrangement of microtubules at the onset of S-phase are unknown. Mammalian CDK1 can phosphorylate β-tubulin and phosphorylated β-tubulin incorporates only poorly into microtubules (Fourest-Lieuvin et al., 2006). Yet, in *Pf*CRK4-depleted parasites, β-tubulin was not differentially phosphorylated (Ganter et al., 2017). A potential other mechanism involves proteins of the kinesin 13 family. These conserved motor proteins possess microtubule depolymerizing activity and function during mitosis in other organisms (Ems-McClung and Walczak, 2010; Moores and Milligan, 2006). *P. falciparum* kinesin 13 was less phosphorylated in *Pf*CRK4-depleted parasites (Ganter et al., 2017), but a potential role of kinesin 13 for microtubule rearrangement remains unclear.

Profiling the *Pf*CRK4 proximity proteome identified 44 proteins in close proximity to the kinase, including *Pf*SPB, which may connect the extra- and intranuclear parts of the centriolar plaque (Fig. S4). Three phosphorylations of *Pf*SPB (S68, S1967, S2195) depend on *Pf*CRK4 (Ganter et al., 2017), but their functional significance remains to be tested. In addition, this set of candidates also informed on how *Pf*CRK4 activity may be regulated. Based on the presence of a conserved amino acid motif in *Pf*CRK4, the pleiotropic kinase *Pf*CK2 was previously predicted to phosphorylate *Pf*CRK4 at serine 977 (Pease et al., 2013). *Pf*CK2 is essential for parasite proliferation in the blood stage (Hitz et al., 2021) and we found the catalytic alpha subunit of *Pf*CK2 (PF3D7_1108400) in the proximity of *Pf*CRK4 (Fig. S3, Tab. S1), supporting a role for *Pf*CK2 in regulating *Pf*CRK4 activity. We also found the putative CDK-regulatory subunit (PF3D7_0105800) enriched in the vicinity of *Pf*CRK4 (Fig. S3, Tab. S1), but its potential role remains elusive. Besides trans-acting factors, autophosphorylation likely also plays a role for *Pf*CRK4 activity as the kinase itself is less phosphorylated in *Pf*CRK4-depleted parasites (Ganter et al., 2017).

The asynchrony during nuclear multiplication suggests that the activity of *Pf*CRK4 is regulated at the level of individual nuclei, which is supported by the uneven distribution of *Pf*CRK4 among nuclei as well as inside nuclei (Fig. 2; S1D, S6). In analogy to canonical cell cycle kinases (Basu et al., 2022; Coudreuse and Nurse, 2010; Zegerman, 2015), *Pf*CRK4 activity is likely altered during parts of the nuclear cycle to allow for genome segregation and licensing of origins (Fig. 4D).

More studies are needed to elucidate the regulation of *Pf*CRK4 activity during asynchronous nuclear cycles and to further analyze its role for other stages of the life cycle (Ganter et al., 2017). Depletion of *Pf*CRK4 had no detectable effect on the development of male gametes, which includes DNA replication, but subsequent parasite development in the mosquito was impaired (Ganter et al., 2017). Recent work has shown that DNA replication and possibly nuclear division during male gamete development is controlled by the related kinase CRK5 (Balestra et al., 2020). Interestingly, CRK5 appears not essential for proliferation in erythrocytes (Balestra et al., 2020; Dorin-Semblat et al., 2013), but if and how both kinases interact is unknown.

Key transitions of the *P. falciparum* nuclear cycle, such es entry into S-phase, are likely under tight control. In absence of canonical cell cycle checkpoints in *P. falciparum* (Matthews et al., 2018), linking entry into S-phase and the initiation of mitotic events via *Pf*CRK4 may be an alternative means to exert control over the nuclear cycle and, in turn, parasite proliferation. In addition, this work renders *Pf*CRK4 a major regulator of nuclear cycle progression and further support its significance as target for chemotherapeutic intervention to curb malaria.

## Material and Methods

### *P. falciparum* cell culture

*P. falciparum* 3D7 parasites were grown in fresh O, Rh+ erythrocytes at 4% hematocrit in RPMI 1640 medium supplemented with 0.5% AlbuMAX II (Gibco), 0.2 mM hypoxanthine (CCPro), 25 mM HEPES, pH 7.4 (Sigma), and 12.5 μg/ml gentamicin (Carl Roth), at 37 °C in 90% relative humidity, 5% O_2_, and 3% CO_2_ (Trager and Jensen, 1976). Routine synchronizations were performed by 5% sorbitol treatment as previously described (Lambros and Vanderberg, 1979). For tight synchronization (1-3 hours), a combination of sorbitol and heparin (50 U ml^-1^) was used (Boyle et al., 2010). Briefly, trophozoite cultures were grown in the presence of 50 IU heparin (Sigma) until most parasites were at the late schizont stage. Heparin was then washed out and parasites allowed to reinvade into fresh RBC for 1 to 3 hours with gentle agitation at 60 rpm. After the invasion period, a sorbitol treatment was performed to remove unruptured schizonts. Parasite genomic DNA extraction was performed on 100-200 μl washed erythrocyte pellet with a parasitemia of 1-5%, using the DNeasy blood & tissue kit (Qiagen).

### Cloning of DNA constructs for parasite transfection

The C-terminus of *Pf*CRK4 (PF3D7_0317200) was tagged with BirA* or GFP by single crossover homologous recombination using the selection-linked integration (SLI) strategy (Birnbaum et al., 2017). For the *Pf*CRK4::BirA* construct, we first amplified the BirA* sequence from MDV1/PEG3::BirA* (Kehrer et al., 2016), using primers P1/P2 and cloned into a pSLI-TGD-based plasmid (a kind gift from Tobias Spielmann) using Mlu1 and SalI restriction sites. Subsequently, the C-terminal 800 bp of *Pf*CRK4 (without stop codon) were PCR amplified from *P. falciparum* 3D7 gDNA using primers P3/P4 and ligated into the NotI/MluI digested pSLI-TGD::BirA* plasmid, giving rise to the pSLI-CRK4::BirA* plasmid. For the *Pf*CRK4::GFP plasmid, the same 800 bp homology region of *Pf*CRK4 (P3/P5) was cloned into pSLI-GFP-glmS (derived from pSLI-TGD) using the NotI/MluI digested backbone and Gibson assembled.

Diagnostic PCR was performed to test for integration of the BirA* (5’ integration: P6/P7, 3’ integration: P8/P9, WT locus: P6/P9) and GFP (5’ integration: P6/P10, 3’ integration: P8/P9, WT locus: P6/P9) targeting constructs. A schematic representation of endogenous *Pf*CRK4, the constructs, and the recombined locus is shown in Fig. S1.

Endogenous tagging of *Pf*SPB (PF3D7_0303500) with a Halo tag was performed by CRISPR/Cas9-mediated homologous recombination using the pDC2-cam-coCas9-U6.2-Hdhfr plasmid (kind gift from Marcus Lee) following previously described methodology (Lee et al., 2019; Lim et al., 2016) and schematically shown in Fig. S4A. To tag *Pf*SPB in the *Pf*CRK4::GFP back ground, which expresses the hDHFR cassette, we first modified the pDC2-cam-coCas9-U6-hDHFR plasmid by replacing the hDHFR resistance marker for a blasticidin (BSD) resistance cassette, producing the construct pDC2-cam-coCas9-U6-BSD. The sequence of the BSD resistance cassette was amplified from the vector p3xNLS-mCherry (Simon et al., 2021).

To remove the BbsI cut site within the BSD gene, which would interfere with the insertion of the gRNAs, we introduced a silent mutation to destroy the BbsI restriction site. For this, the BSD cassette was amplified as two fragments overlapping at the restriction site (P11/P12 and P13/P14), with the mutation introduced in the respective overlapping forward and reverse primers and cloned into the EcoRI-ApaI sites of the vector, ultimately creating the vector pDC2-cam-coCas9-U6.2-BSD. The selection of gRNA sequences, designed using the Eukaryotic Pathogen CRISPR guide RNA/DNA Design Tool (Peng and Tarleton, 2015), was based on the distance to the C-terminus of *Pf*SPB and the predicted activity scores. gRNAs carrying overhangs complementary to the BbsI cut site (P15/P16) were annealed and phosphorylated before ligation into the BbsI digested pDC2-cam-coCas9-U6.2-BSD plasmid.

Next, the left and right homologous arms (donor template) were amplified separately by PCR from genomic DNA of *P. falciparum* 3D7. In total four overlapping fragments were amplified: the 5’ homology region as two parts to include protective silent mutations, which avoid recutting after recombination (P17/18 and P23/P24), the 3’ homology region (P21/P22), as well as the Halo tag sequence with a stop codon (P19/P20). These fragments where Gibson assembled and ligated into the EcoRI and AatII digested plasmid (Fig. S4A).

All primer sequences used to generate and genotype the mutant parasite lines are listed in Tab. S1. Primer were purchased from Thermo Fisher Scientific or Sigma Aldrich. Restriction enzymes were purchased from NEB. All PCRs were performed using Phusion DNA polymerase (NEB). Molecular cloning was performed using HiFi DNA Assembly (NEB) T4 DNA Ligase (NEB) or Gibson Assembly (NEB). Correct sequence of inserts was verified by Sanger sequencing.

### Transfection of *P. falciparum*

For transfection, young ring stage parasites at 5-8% parasitemia were transfected with 75-100 μg of purified plasmid DNA as previously described (Fidock and Wellems, 1997). Electroporation was performed with a Gene Pulser Xcell, Bio-Rad (Settings: 0.31 kV, 0.950 μF, capacitance set to “High Cap”, resistance on the Pulse Controller II set to “Infinite”). To select for parasites carrying the genomic modification via the SLI system, we followed the previously published protocol (Birnbaum et al., 2017). In brief, a first selection for episomally maintained plasmids was carried out using 2.5 nM of WR99210 (kind gift of Jacobus Pharmaceutical Company) followed by treatment with 800 μg/ml geneticin (Thermo Fisher Scientific) for genome integration selection. Limiting dilution was done to obtain clonal parasite lines. For the CRISPR/Cas9 approach, selection was performed with 5 μg/ml blasticidin S (InVivoGen). Unless otherwise noted, *Pf*CRK4–HA–DD parasites were cultured in the presence of 250 nM Shield-1 (Ganter et al., 2017).

### Live and fixed cell imaging

#### Immunofluorescence assay

Immunofluorescence staining for confocal microscopy of bloodstage parasites was done as previously described with minor alterations (Simon et al., 2021). In brief, parasitized erythrocytes were seeded on poly-L-lysine coated ibidi 8-well chambered glass bottom dish and fixed with 4% PFA/PBS at 37°C for 20 min. Cells were then permeabilized with 0.1% Triton X-100/PBS for 15 min at room temperature, rinsed three times with PBS and incubated with 0.1 mg/ml NaBH_4_/PBS solution for 10 min to quench free aldehyde groups. After washing thrice with PBS, cells where blocked with 3% BSA/PBS for 30-60 min and incubated with primary antibodies diluted in 3% BSA/PBS for 2 h at room temperature or overnight at 4°C. Next, the cells were washed three times with 0.5% Tween-20/PBS and incubated with secondary antibodies plus Hoechst in 3% BSA/PBS for 1 h at room temperature. After washing twice with 0.5% Tween-20/PBS and once with PBS, cells were directly imaged or stored in PBS at 4°C in the dark until imaging.

For the *Pf*SPB::Halo parasites line, labelling was performed in live cells accordingly to the manufacturer instructions (Promega). In brief, parasites were labelled for 1 hour at 37°C in standard culture conditions with 5 μM of the cell permeable HaloTag^®^ TMR ligand. After labelling, cells were rinsed with fresh prewarmed imaging media for 30 minutes, fixed and processed for immunofluorescent staining as described above.

Images acquisition was done using a Leica TCS Sp8 scanning confocal microscope using the HC PL APO CS2 63x/1.4 N.A. oil immersion objective. Unless otherwise stated, images were captured as a series z-stacks separated by 0.2-0.4 μm intervals and representative confocal images are shown as maximum projections. Images were processed with the Lightning algorithm in the adaptive mode (except for Fig. 1, 2, S1) using default settings with further analysis using Fiji (Schindelin et al., 2012). Brightness and contrast were adjusted for visualization.

#### Analysis of the cellular distribution of *Pf*CRK4

To quantify the *Pf*CRK4 distribution relative to microtubules, the signal density of *Pf*CRK4::GFP was determined in different cellular compartments and at different microtubule structures (Fig. 2; S6, S7). These areas were defined by the presence of different markers. Specifically, the nuclear compartment was defined as the area with a high Hoechst 33342 signal and microtubules as the areas with a high α-tubulin antibody signal. The whole parasite (including the nucleus but without the food vacuole) was defined as the area where there was weak, unspecific background signal of the α-tubulin antibody, which was only present in the parasite but not the red blood cell, seen by comparison with the brightfield channel. The cytoplasm compartment was set as the whole parasite (as defined above) without the areas that showed a high Hoechst 33342 signal, i.e., the nuclei.

The different areas were isolated by creation of specific masks from the reference channels through thresholding. Before thresholding, a gaussian blur with a radius of 2 pixel was applied to the reference channels. Subsequently, the reference channels were thresholded in the default mode with the following parameters: nuclear mask (Hoechst 33342 channel), threshold of 30/255; parasite mask (α-tubulin channel), threshold of 10/255; microtubule mask (α-tubulin channel), threshold of 50/255. For the cytoplasm mask (parasite mask minus the nucleus mask), the area of the nucleus mask was removed from the corresponding parasite mask by inverting the nucleus mask and multiplying it with the cytoplasm mask via the image calculator function of Fiji.

The reference masks were then individually applied to the unmodified *Pf*CRK4 channel and all *Pf*CRK4 signal outside of the respective mask was deleted. The remaining *Pf*CRK4 signal in the different z-slices was added using the Sum Slices mode of the z-projection function in Fiji and measured as the raw integrated density. To obtain the signal density within the different compartments, the raw integrated density of *Pf*CRK4 within that compartment was divided by the area of the respective mask that was applied. The area of the respective masks was determined by thresholding the binary masks, measuring the area of the threshold and summing the area of the different z-slices. *Pf*CRK4 signal densities within the different compartments were normalized to the signal density within the parasite area.

To stratify the different microtubule structures, the microtubule masks were manually segmented in 3D by removing all areas that did not belong to either an elongated or a focal microtubule signal. Subsequently, these masks containing either those areas of elongated or focal microtubules were used to isolate the total signal of *Pf*CRK4 in only the selected microtubule structures and the signal within these masks measured as detailed above. For comparison of *Pf*CRK4 signal density at different microtubule structures within the same parasite, *Pf*CRK4 signal densities at different microtubule structures were normalized to the mean signal intensity of *Pf*CRK4 at all microtubule structures within the same cell.

#### Live-cell imaging

For live cell imaging, we followed previously published protocols with minor modifications (Klaus et al., 2022). In brief, sterile glass bottom 8 well-dishes (ibidi GmbH) were coated with 5 mg/ml Concanavalin A (Sigma-Aldrich) and rinsed with PBS. 500 μl of resuspended parasite culture was washed twice with incomplete RPMI and left to settle on the dish for 10 min at 37°C before unattached cells were washed off using incomplete RPMI until a monolayer remained. Cells were left to recover at standard culturing conditions in complete RPMI for at least 8 h before media was exchanged to the phenol red-free complete RPMI imaging medium (RPMI 1640 L-Glutamine (PAN-Biotech), 0.5% AlbuMAX II, 0.2 mM Hypoxanthine, 25 mM HEPES pH 7.3, 12.5 μg/ml gentamicin) that had been equilibrated to incubator gas conditions for at least 6 h.

Cells were incubated with SPY555-tubulin (spirochrome) in phenol red-free complete RPMI imaging medium starting from 2 h before start of imaging. For experiments presented in Fig. 3 and movies S1–4, cells were additionally incubated with 5-SiR-Hoechst kindly provided by Gražvydas Lukinavičius and Jonas Bucevičius (Bucevičius et al., 2018). For *Pf*CRK4 rescue experiments, *Pf*CRK4–HA–DD was stabilized at the same time as addition of the live-cell compatible dyes by the addition of 250 nM of Shield-1 within the imaging medium. Dishes were completely filled with imaging media, closed and sealed tightly using parafilm before imaging started.

Point laser scanning confocal microscopy was performed on a Zeiss LSM900 microscope equipped with an Airyscan 2 detector using Plan-Apochromat 63x/1,4 oil immersion objective. Live-cell imaging was performed at 37°C. Images were acquired at multiple positions using an automated stage and the Definite Focus module for focus stabilization with a time-resolution of 15 min/stack for 2-5 h for correlation of microtubule structure with DNA content (Fig. 3; movies S1–4), and with a time-resolution of 10 min/stack for 15 h for correlation of microtubule structures and DNA replication (Fig. 4; movies S5, S6). Multichannel images were acquired sequentially in the line scanning mode using 561 nm and 640 nm diode lasers for SPY555-tubulin and 5-SiR-Hoechst imaging, respectively. Emission detection was configured using variable dichroic mirrors to be 490-650 for SPY555-tubulin and 490-700 for 5-SiR-Hoechst detection. The Airyscan detector was used with the gain adjusted between 700 and 900 V, offset was not adjusted (0%). Brightfield images were obtained from a transmitted light PMT detector, with the gain adjusted between 700 and 900 V. Sampling was Nyquist-optimized in xy axis (approx. 50 nm) and 500 μm in z axis, bidirectionally with pixel dwell time between 0.7 and 1.2 μs and 2x line averaging. Subsequently, ZEN Blue 3.1 software was used for 3D Airyscan processing with automatically determined default Airyscan Filtering (AF) strength.

#### Analysis of live-cell imaging to correlate microtubule structures with DNA content

Processing and analysis of imaging files was done with Fiji (Schindelin et al., 2012), analysis and plotting of data in Microsoft Excel and Prism GraphPad 8, respectively. Multiposition images were manually inspected for dead or abnormal parasites using the brightfield channel. Individual parasites were saved as 300 x 300-pixel TIFF files containing original channel, time and z-slice information. Parasites were further evaluated for nuclei number and occurrence of hemispindle structures. Only parasites containing one nucleus and distinct hemispindle structures were chosen for further analysis.

Time-lapse images were stabilized in xy using the Register Virtual Stack Slices Plugin in Fiji. Briefly, for each parasite a time-lapsed reference z-slice of the brightfield channel was registered via the translation mode (no deformation) at standard parameters. The resulting transformation matrixes were saved. The transformation matrixes were then applied to all other z-slices in all other channels using the Transform Virtual Stack Slices Plugin in Fiji.

For analysis of DNA content and microtubule structures (Fig. 3C–F), a tight selection was first applied around the parasite to remove unspecific background signal outside of the infected red blood cell, which was particularly strong in the 5-SiR-Hoechst channel. For this, an oval mask of varying size was applied around the parasite in the 5-SiR-Hoechst channel and all signal outside of the mask deleted. To further subtract background and segment the nucleus a second mask was applied, which was created from the background-subtracted 5-SiR-Hoechst channel by applying a median filter with a radius of 2 pixels and a threshold of 550 or 300 depending on signal intensity. The resulting mask was applied to the 5-SiR-Hoechst channel, all signal outside the mask deleted, the signal in the different z-slices of the resulting image summed using the z-projection Sum Slices function and the remaining raw integrated density measured. For analysis of the DNA content over time, the raw integrated density was normalized to the first timepoint of the timelapse of the individual parasites.

For visualization of the different microtubule structures, a collage of the SPY555-tubulin channel over time was done. A field of 100 x 100 pixels centered around the spindle structures in a maximum intensity z-projection of the SPY555-tubulin channel was cut out and the resulting images combined in a single row via the make montage function in Fiji without a border. All antibodies and dyes used in this study are detailed in Tab. S2.

### Cell culture and harvest of biotinylated proteins

Experiments were performed as described previously with minor modifications (Boucher et al., 2018; Khosh-Naucke et al., 2018; Roux et al., 2012). To label biotinylated proteins in parasites for analysis by fixed imaging, streptavidin blot, or mass spectrometry, sorbitol synchronized ring stage *Pf*CRK4::BirA* and 3D7 WT parasites were cultured [+] or [−] of 50 uM biotin (Sigma) supplementation. Cultures were harvested for analysis 24 hours later as the majority of parasites had multiple nuclei.

For mass spectrometry identification of biotinylated proteins, ~1× 10^9^ parasites (4 mL of packed red blood cells with 8-10% parasitemia) were isolated in triplicate for each condition. Uninfected erythrocytes were removed using saponin buffer (0,1% saponin/PBS plus 1× cOmplete™ protease inhibitor cocktail (Roche)). Cells were washed twice with saponin buffer to further remove erythrocytes and erythrocyte debris, followed by PBS plus 1× cOmplete™ protease inhibitor cocktail and 1 mM phenylmethylsulfonyl fluoride. Parasite pellets were resuspended in 1 mL of RIPA lysis buffer (50 mM Tris, 150 mM NaCl, 0.1% SDS, 0.5% sodium deoxychlorate, 1% NP-40, 1 mM EDTA, 1 mM dithiothreitol (DTT) and 1× cOmplete™ protease inhibitor) for 15 minutes and sonicated in a water bath Bioruptor Plus (10 cycles of 30 sec ON and 30 sec OFF pulses at high intensity). The lysates were further clarified by centrifuging at 15000 g for 15 min at 4°C to remove hemozoin and other insoluble debris. 100 μl of streptavidin magnetic beads (MagResyn Streptavidin beads) were washed twice with 0,1% BSA in PBS and blocked with 2.5% BSA in PBS for 1 h at room temperature with gentile agitation on a rotating wheel. Beads were washed twice with RIPA buffer and incubated with 1 ml of each sample lysate over night at 4°C with gentile agitation. After incubation, beads were sequentially washed thrice with RIPA buffer, thrice with SDS buffer (2% SDS, 50 mM Tris-HCl, pH 7.4, 150 mM NaCl), thrice with Urea buffer (8 M Urea, 50 mM Tris-HCl, pH 7.4, 150 mM NaCl) and resuspended in 400 μl of 100 mM ammonium bicarbonate buffer, pH 8.0, and stored at −80°C until further processing. A fraction of the beads samples was taken for Western blot analysis.

### SDS Page and Western Blotting

Samples for Western blot analysis were supplemented with 4X Laemmli buffer (Bio-Rad) and 2.5%β-mercaptoethanol. After incubation for 10 min at 95 °C, beads were removed via a magnetic rack and proteins loaded on a 4–15% Mini-PROTEAN^®^ TGX™ Precast Protein Gel (Bio-Rad). Blotting was performed using the Trans-Blot Turbo Mini 0.2 μm Nitrocellulose Transfer Pack (Bio-Rad) on a Trans-Blot Turbo Transfer System (Bio-Rad). The membrane was blocked with 3% BSA in TBST (0.05% Tween-20/PBS) for 1 hour and then probed with α-histone H3 antibody in 1% BSA/TBST for 1 hour at room temperature. After three washes with TBST for 10 min, the membrane was stained with the secondary antibodies IRDye 800CW-labeled streptavidin and IRDye 680CW Goat anti-rabbit IgG in 1% BSA/TBST supplemented with 0.05% SDS for 1 hour at room temperature (Tab. S2). After three 10 min washes with TBST and one 5 min wash with PBS, fluorescent signals were visualized using a LI-COR Odyssey^®^ CLx Imaging System.

### Label-free quantitative mass spectrometric analysis

Label-free quantitative mass spectrometry and data analysis were performed as commercially available service by the Proteome Factory, Berlin, Germany (www.proteome-factory.com) according to standard procedures.

#### On-Beads Trypsin Digestion

Proteins bound to magnetic beads were cleaved by trypsin according to standard procedures. In brief, magnetic beads were washed twice with 45 μl ABC buffer (ammonium bicarbonate buffer, 50 mM, pH 8.0), then resuspended in further 45 μl ABC. Reduction of cysteine residues was performed by addition of 5 μl 100 mM DTT and incubation for one hour at 60°C. For alkylation, 6 μl 100 mM 2-iodoacetamide were added and incubated for one hour at room temperature. Residual alkylating agent was quenched by further addition of 100 mM DTT. 500 ng Trypsin was added, and proteolysis allowed to proceed overnight at 37°C. The solution was acidified by addition of formic acid and the supernatant collected. Samples were stored at −80°C until analysis.

#### LC-MS/MS Analysis

Label-free quantitative proteomics using LCMS/MS was performed using an Ultimate 3000 nanoHPLC system. Samples were loaded onto a trapping column (kept at 35°C) for desalting (Thermo Fisher Scientific, Dionex, Pepmap C18) and peptides separated on a 500 × 0.075 mm column (0.5 μl/min, 50°C, Reprosil C18-AQ, Dr. Maisch) using a linear gradient from 10% to 32% B (solvent A: water, solvent B: acetonitrile, both with 0.1% formic acid). The column effluent was directed to an Orbitrap Velos mass spectrometer via a nanoeletrospray ion source. Survey scans were detected at a nominal resolution of R=60,000, and up to ten MS/MS spectra from ions of interest (charge states +2 and above) were data-dependently recorded. Total acquisition time was 300 min per analysis.

#### Database search and analysis

RAW data were processed with the MaxQuant software package (version 1.6.14.0). MSMS spectra were searched against *P. falciparum* 3D7 and human sequences from RefSeq databases. Cysteine carbamidomethylation was set as a fixed modification while methionine oxidation and glutamine (Gln) and asparagine (Asn) deamidation were used as variable modifications for database search. Label-free quantification was done with oxidation (M), acetylation (Protein N-term) and deamidation (N) included and using the match between runs feature. Default settings were used for further parameters.

Statistical analysis of the proteingroups.txt file from the MaxQuant output was done with Perseus software (version 1.6.15.0). LFQ data was imported, then decoy matches and putative contaminants removed. Intensity data was log2 transformed, and proteins with at least two valid LFQ values in each group were further analyzed. Imputation using the “imputeLCMD” tool from the PerseusR Package was used with MCARS and MNARS+ where applicable. Student’s t-test was applied to resulting data and a significance threshold set to *P*-value < 0.05. Fold Change was calculated from the difference of the mean LFQ values from *Pf*CRK4::BirA* [+] biotin and *Pf*CRK4::BirA* [−] biotin by retransforming log2 values. For negative log2 fold changes, the negative inverse value of the 2^x value was given.

Applying correction for multiple hypothesis testing using Benjamini-Hochberg or permutation based FDR resulted in no significantly different protein groups between *Pf*CRK4::BirA* [+] and [–] biotin.

#### Gene Ontology (GO) term enrichment analysis

GO term enrichment analysis was performed for molecular function, biological process, and cellular component of both overrepresented and underrepresented proteins using PlasmoDB (www.plasmodb.org, accessed on 2 Abril 2022). The following parameters were used: evidence: curated; limit to GO slim terms: yes; *P*-value from Fisher’s exact test ≤ 0.05.

#### STRING analysis

Interaction evidence among the overrepresented proteins in the vicinity of *Pf*CRK4 was analyzed using the STRING database, version 11.5 (www.string-db.org, accessed on 2 August 2022) (Szklarczyk et al., 2020). In Fig. S5B proteins are presented as nodes (circles) connected by lines (Edge), whose thickness represents the strength of the connection based on the STRING database. For simplicity, only direct interactions with *Pf*CRK4 are shown in Fig. S5B. The entire dataset is provided in Tab. S3. The functional linkages predicted here were based on experiments, neighborhood, fusion-fission events, co-occurrence and coexpression, and data imported from public databases of physical interactions. A medium confidence cut-off of 0.4 was used.

### Flow cytometry analysis of parasite DNA content and growth rate

Parasites DNA content and growth rate were determined as previously described using SYBR^®^ Green I (Ganter et al., 2017; Russo et al., 2009; Theron et al., 2010). In brief, synchronized (± 1 h) *Pf*CRK4–HA–DD parasites were grown in the presence or absence of 250 nM of Shield-1(Fig. S8A). At 32 hpi, 500 nM of Paclitaxel (PTX) or solvent was added to the cultures. At different times points (Fig. S8C, D), 100 μl of resuspended culture was collected and fixed with 4% PFA/PBS and 0.0075% glutaraldehyde for 30 min at room temperature. To reduce RNA-derived background signal, cells were permeabilized with Triton™ X–100 (Sigma, T8787) for 8 min, treated with 0.3 mg / ml Ribonuclease A (Sigma, R4642) for 30 min and subsequently washed. Cells were stained for 20 min with 1:2000 SYBR^®^ green I (Invitrogen) in PBS, washed, and analyzed on a BD FACSCelesta flow cytometer. For all conditions, 500-1000 infected red blood cells were recorded. To calculate cellular DNA content, fluorescence measurements from single-infected parasites at 24 hpi were defined as 1 *C*. All flow-cytometry data were analyzed with FlowJo and GraphPad software.

## Acknowledgements

The authors thank F. Hentzschel for critically reading the manuscript and members of the Ganter and Guizetti groups for ongoing discussions. We thank MT. Duraisingh for kindly providing the *Pf*CRK4–HA–DD parasite line, G. Lukinavičius and J. Bucevičius for providing the 5’-SiR-Hoechst dye, T. Spielman for the pSLI-TGD plasmids, M. Lee for the CRISPR-Cas9 plasmid, and Jacobus Pharmaceutical Company for WR99210. We are grateful to the Infectious Diseases Imaging Platform (IDIP) at the Center for Integrative Infectious Disease Research, Heidelberg, Germany, for the microscopy support. The *Plasmodium* database PlasmoDB (www.plasmodb.org) greatly facilitated this work.

## Additional information

### Funding

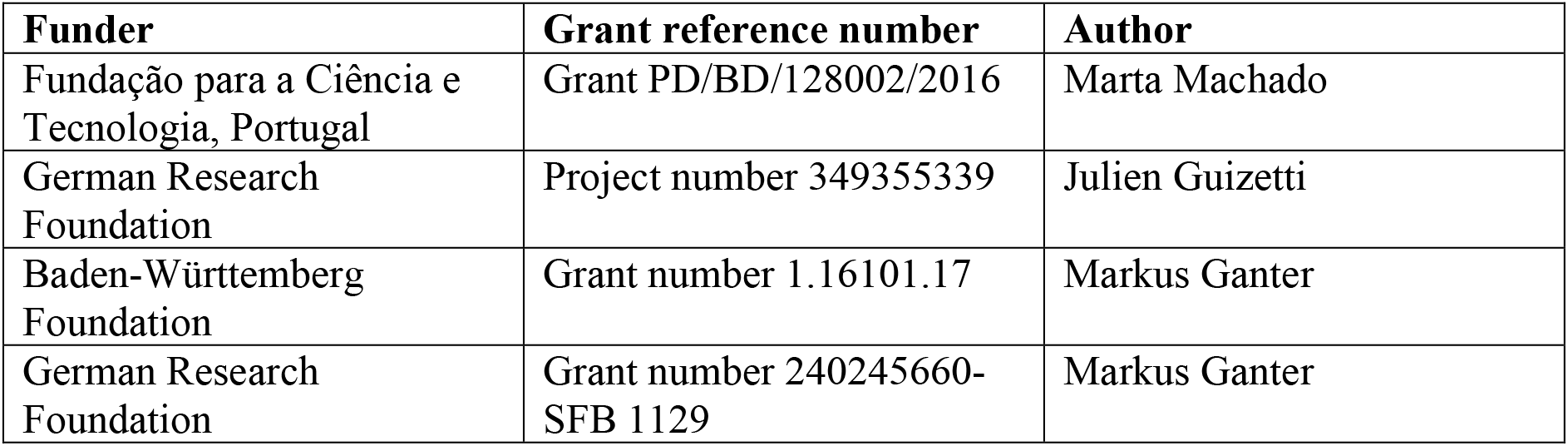

The funders had no role in study design; data collection, analysis, and interpretation; or publication of this work.

### Author contribution

Conceptualization: M.M., S.K., and M.G.; Methodology: M.M., S.K., and M.G.; Software: S.K.; Investigation: M.M. and S.K.; Formal Analysis: M.M. and S.K.; Validation: J.G.; Resources: D.K.; Writing—original draft: M.M. and M.G.; Writing—review and editing: M.M, S.K., and M.G.; Visualization: M.M, S.K., and M.G.; Funding acquisition: M.M. and M.G.; Supervision: M.G.

## Additional files

### Supplementary files

**Fig. S1.**
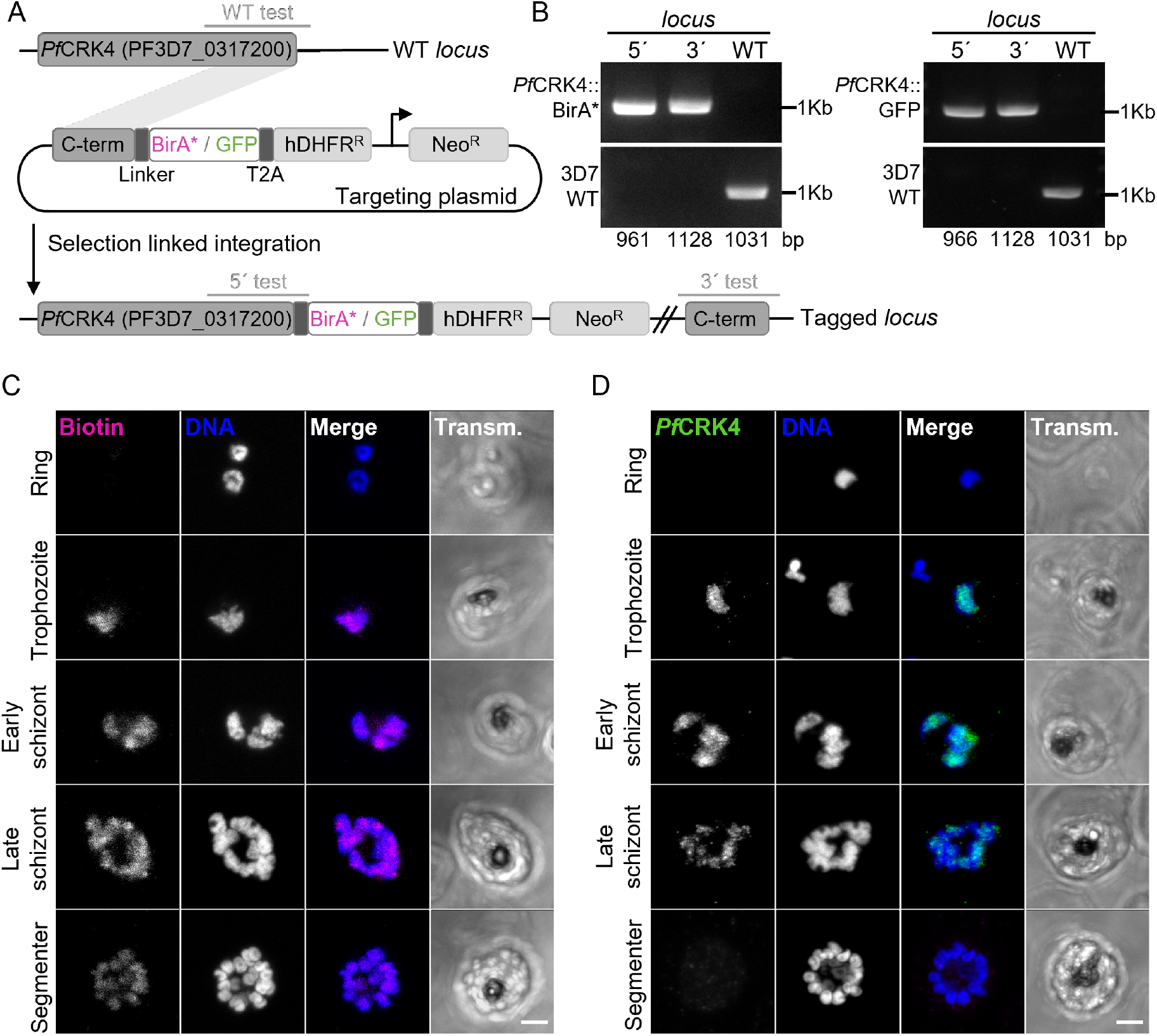
Expression of *Pf*CRK4::BirA* leads to biotinylation of proteins largely restricted to the nucleus and its periphery, consistent with the localization of *Pf*CRK4. **A** Schematic representation of the SLI-based strategy used to fuse endogenous *Pf*CRK4 with BirA* or GFP. This vector integrates into the C-terminal part of *Pf*CRK4 via single-crossover recombination and, only upon integration conveys resistance to geneticin. The expression of BirA* or GFP is driven by the endogenous *Pf*CRK4 promoter. Expected products for diagnostic PCR are shown as grey lines. The shown linker was not added to the *Pf*CRK4::BirA* construct. **B** Diagnostic PCR using genomic DNA from clonal *Pf*CRK4::BirA* (left panel) or GFP lines (right panel) and *P. falciparum* 3D7 wild type (WT) parasites showing successful plasmid integration. All primer sequences are listed in Tab. S2. **C** Representative confocal images of *Pf*CRK4::BirA* parasites grown for 24h in media supplemented with 50 μM of biotin. Biotinylated proteins were detected with Atto647N-labeled streptavidin (magenta). **D** Representative confocal images showing parasites endogenously expressing GFP-tagged *Pf*CRK4 labelled with anti-GFP (green). Scale bars 2 μm.

**Fig. S2.**
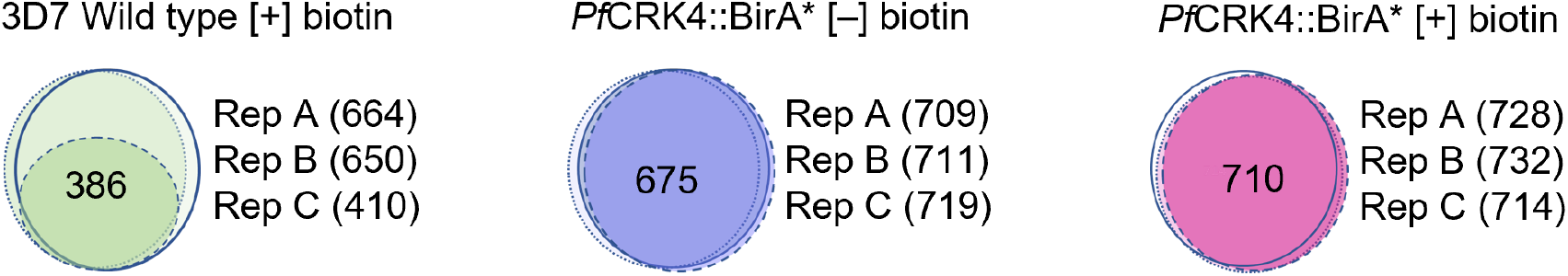
Venn diagrams of identified proteins in three biological replicates. Shown are the overlap of all proteins identified with at least two spectral counts by mass spectrometry in three biological replicates of each condition tested: 3D7 Wild type [+] biotin, *Pf*CRK4::BirA* [−] biotin and *Pf*CRK4::BirA* [+] biotin.

**Fig. S3.**
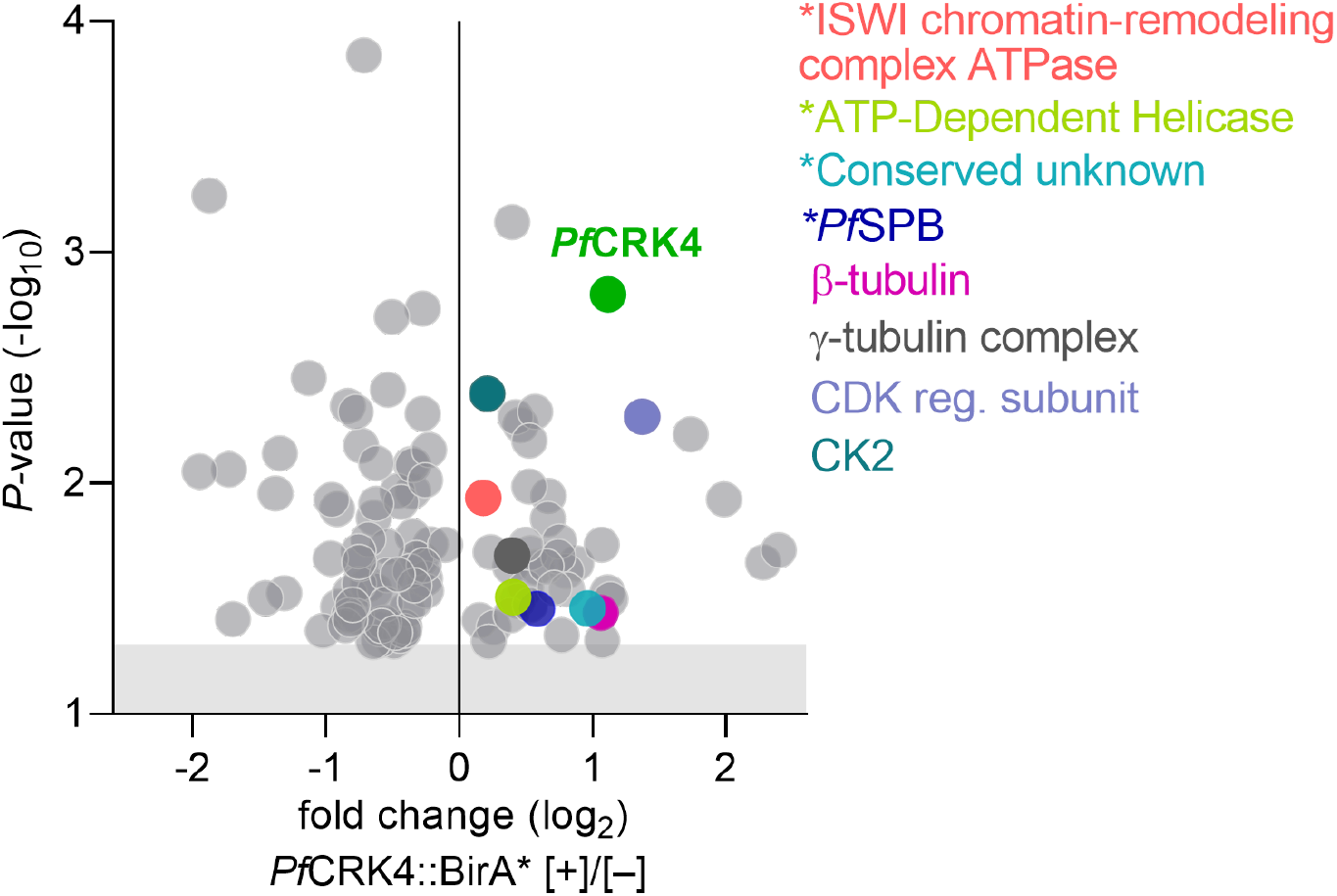
Under- and overrepresented proteins in the vicinity of *pf*CRK4::BirA* parasites [+] 50 μM biotin relative to [−] biotin. Highlighted are the bait protein *Pf*CRK4, tree proteins associated to microtubules (*PF*SPB, β-tubulin, and γ-tubulin complex component), two potential regulators of *Pf*CRK4 activity (CK2 and CDK-regulatory subunit), as well as shared hits between the *Pf*CRK4 proximity proteome and the *Pf*CRK4 phosphoproteome (marked with an asterisk). Shown are mean values of triplicates; grey shaded region indicates a *P*-value ≥ 0.05 (Student’s *t*-test, two-tailed, equal variance).

**Fig. S4.**
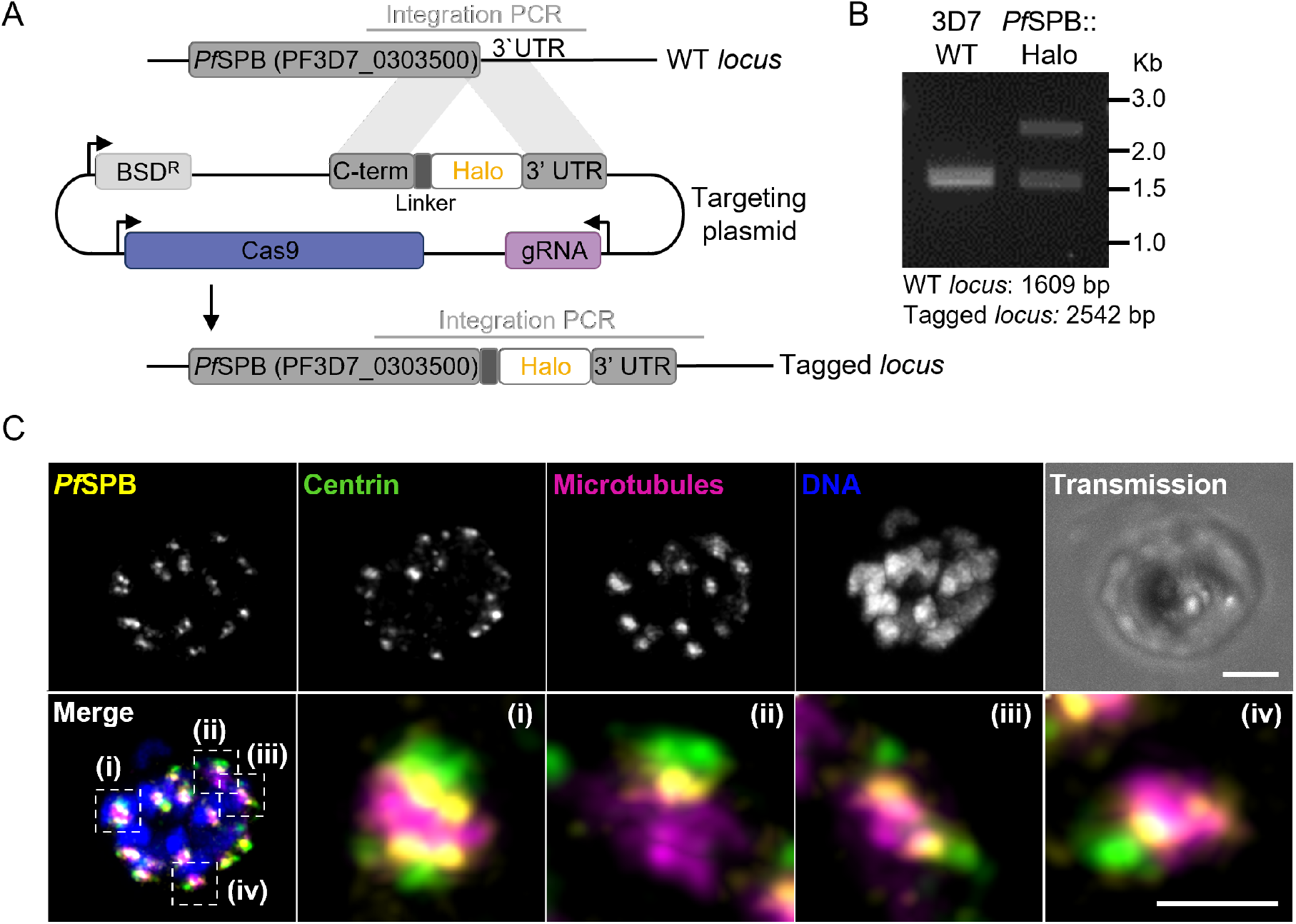
*Pf*SPB localizes to the centriolar plaque. **A** Schematic representation of the CRISPR/Cas9-based strategy used to tag the endogenous candidate protein PF3D7_0303500, a putative spindle pole body protein (*Pf*SPB), with a Halo Tag. Expected products for diagnostic PCR are shown as grey lines. **B** Diagnostic PCR using genomic DNA from a non-clonal *Pf*SPB::Halo and *P. falciparum* 3D7 wild type (WT) parasites showing successful tagging. All primer sequences are listed in Tab. S2. **C** Confocal microscopy showing the cellular localization of *Pf*SPB. Co-labelling with anti-centrin and anti-tubulin antibodies shows that *Pf*SPB localizes to the centriolar plaque, between the intranuclear tubulin and the extranuclear centrin; scale bar, 2 μm, zoom-in 1 μm.

**Fig. S5.**
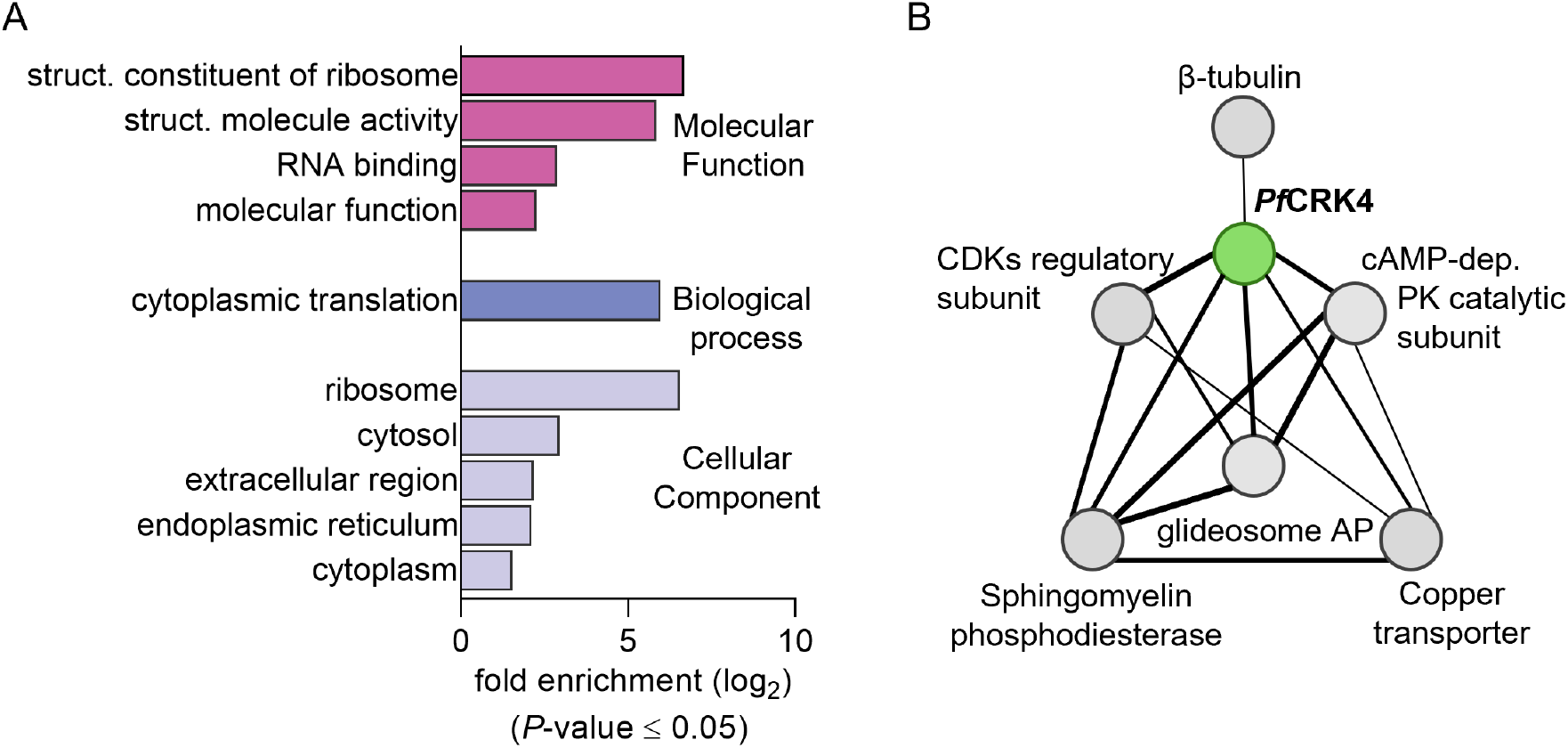
Profiling the proximity proteome of *Pf*CRK4. **A** Gene ontology term enrichment analysis of 71 proteins that are underrepresented in the vicinity of *Pf*CRK4, calculated using the built-in tool at PlasmoDB.org. **B** Protein-protein interaction network of the *Pf*CRK4 interactome adapted from the STRING database. For simplicity, only direct interactions with *Pf*CRK4 are shown. The thickness of the lines indicates the degree of confidence prediction of the interaction (see also Tab. S3).

**Fig. S6.**
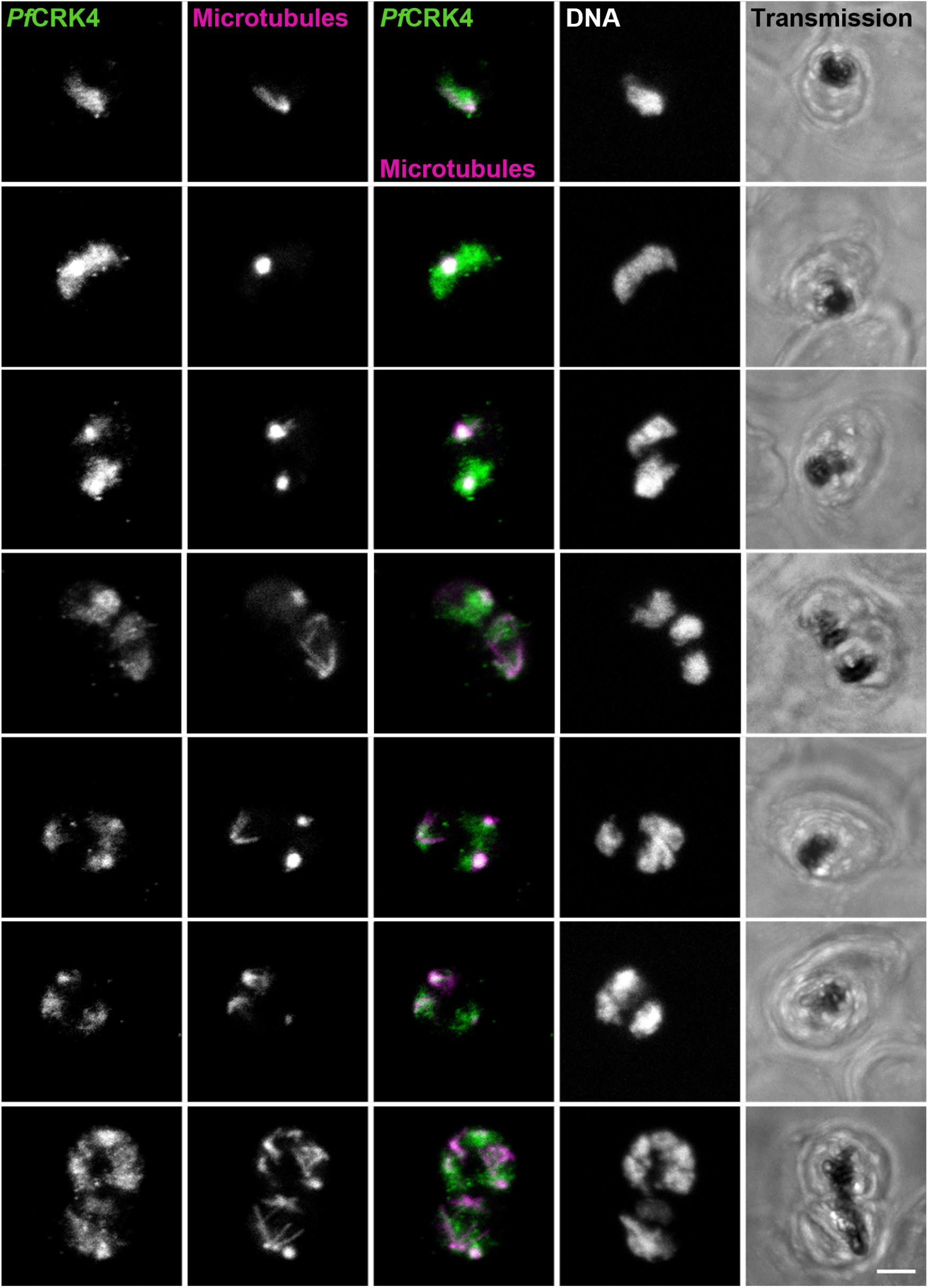
Distribution of *PfCRK4* relative to nuclear microtubule structures in *P. falciparum* blood stages. Immunofluorescence detecting *Pf*CRK4::GFP, microtubules, and DNA; scale bar, 2 μm.

**Fig. S7.**
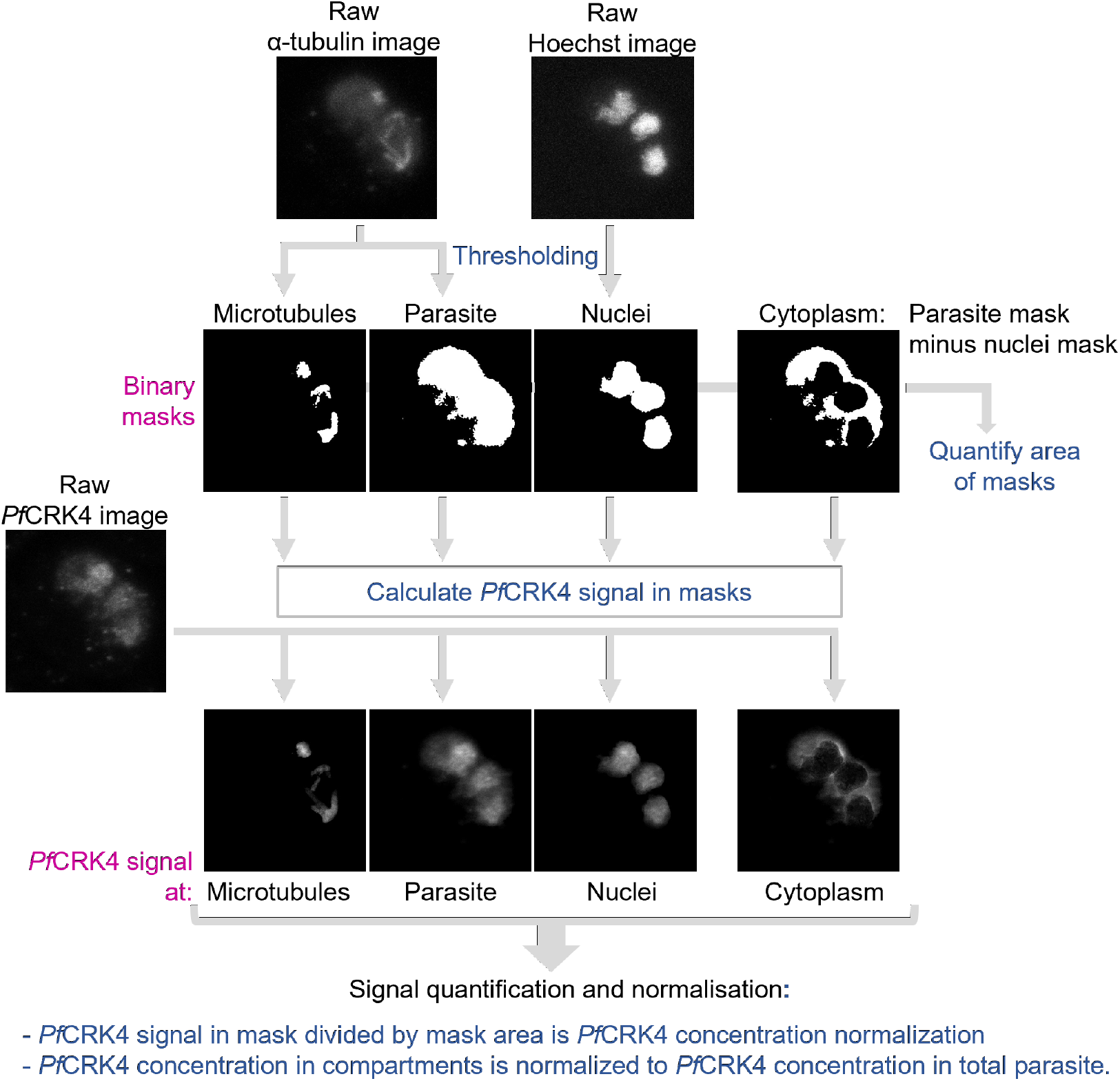
Image analysis workflow to quantify the distribution of *Pf*CRK4 relative to nuclear microtubule structures. A detailed description of the workflow is provided in the Material and Methods section.

**Fig. S8.**
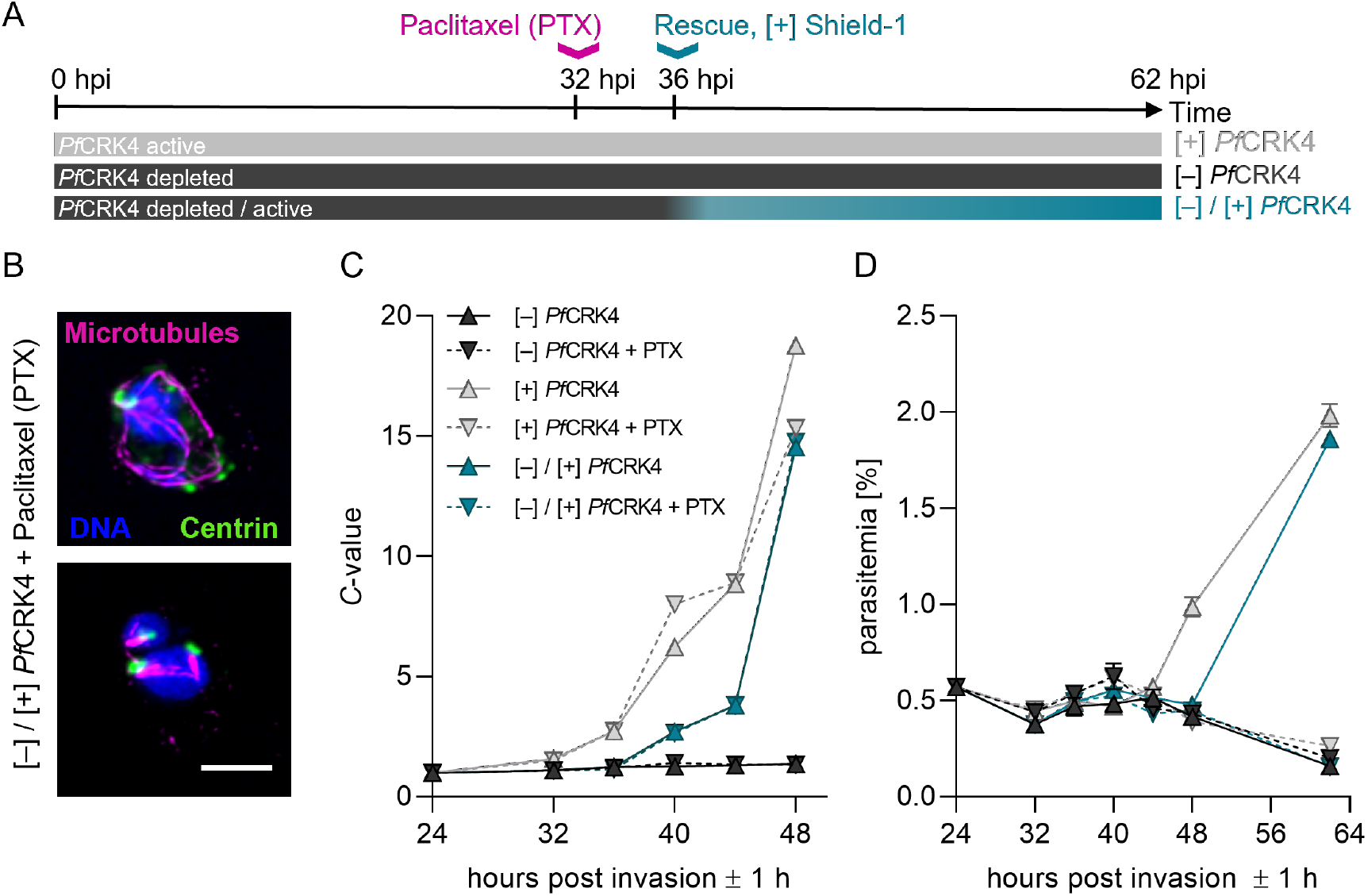
Normal microtubule rearrangement is not a prerequisite for DNA replication. **A** Schematic representation of the experimental workflow, illustrating the timing of paclitaxel (PTX) treatment and *Pf*CRK4 activity; hpi, hours post invasion. **B** Confocal images of two PTX-treated and *Pf*CRK4-depleted parasites with rescued kinase expression from 36 hpi onwards and imaged at 40 hpi; scale bar, 2 μm. **C** DNA content of *Pf*CRK4 parasites treated with PTX or solvent; mean ± SD of triplicates (representative of two biological replicates); ring stage DNA content defined as 1 *C*. **D** Proliferation of *Pf*CRK4 parasites treated with PTX or solvent; mean ± SD of triplicates (representative of two biological replicates).

**Tab. S1.** Proteins identified by label-free quantitative mass spectrometry in three biological replicates of each condition (*P*-value ≤ 0.05).

**Tab. S2.** List of primers, antibodies and dyes.

**Tab. S3.** Interaction network of potential *Pf*CRK4 interactors identified by BioID based on STRING analysis (Szklarczyk et al., 2020).

**Movie S1, S2.** Time-lapse super-resolution microscopy of *Pf*CRK4-depleted parasites showing highly dynamic hemispindle-like structures. Microtubules were labelled with the live-cell compatible dye SPY555-Tubulin (magenta) and DNA with 5-SiR-Hoechst (blue); Transmission image, single slice; fluorescent images, average intensity z-projections; brightness and contrast adjusted for better visibility; time interval, 15 min; scale bar, 5 μm.

**Movie S3, S4.** Time-lapse super-resolution microscopy of parasites with rescued *Pf*CRK4 activity showing rearrangement of microtubules upon restored kinase expression. Imaging conditions as in movies S1, S2; scale bar, 5 μm.

**Movie S5, S6.** Time-lapse super-resolution microscopy of a reporter parasite ectopically expressing nuclear mCherry (blue) and PCNA1::GFP (green) and labelled with SPY555-Tubulin (magenta) showing the dynamics of DNA replication and microtubules organization; Transmission image, single slice; fluorescent images, average intensity z-projections; brightness and contrast adjusted for better visibility; time interval, 10 min; scale bar, 5 μm.

### Data availability

All data needed to evaluate the conclusions in the paper are present in the paper and/or the Supplementary Materials.

